# Necessary Conditions for Reliable Representation of Asynchronous Spikes Through a Single-layered Feedforward Network

**DOI:** 10.1101/538736

**Authors:** Milad Lankarany

## Abstract

Reliable propagation of firing rate – specifically slow modulation of asynchronous spikes in fairly short time windows [20-500]^ms^ across *multiple layers of a feedforward network* (FFN) receiving background synaptic noise has proven difficult to capture in spiking models. We, in this paper, explore how information of asynchronous spikes disrupted in the *first layer of a typical FFN*, and which factors can enable reliable information representation. Our rationale is that the reliable propagation of information across layers of a FFN is likely if that information can be preserved in the first layer of the FFN. In a typical FFN, each layer comprises a certain number (network size) of excitatory neurons – leaky integrate and fire (LIF) model neuron in this paper – receiving correlated input (common stimulus from the upstream layer) plus independent background synaptic noise. We develop a reduced network model of FFN which captures main features of a conventional all-to-all connected FFN. Exploiting the reduced network model, synaptic weights are calculated using a closed-form optimization framework that minimizes the mean squared error between reconstructed stimulus (by spikes of the first layer of FFN) and the original common stimulus. We further explore how representation of asynchronous spikes in a FFN changes with respect to other factors like the network size and the level of background synaptic noise while synaptic weights are optimized for each scenario. We show that not only synaptic weights but also the network size and the level of background synaptic noise are crucial to preserve a reliable representation of asynchronous spikes in the first layer of a FFN. This work sheds light in better understanding of how information of slowly time-varying fluctuations of the firing rate can be transmitted in multi-layered FFNs.

## I. Introduction

The brain is highly modular. Effective communication between modules relies on the reliable transmission of information. Feedforward connections are responsible for linking upstream neurons with downstream neurons, either across different layers within the same cortical region, or between different cortical regions. Notwithstanding the influence of lateral and recurrent connections, multi-layered feedforward neural networks (FFN) play a critical role in conveying information within the brain [1].

Information can be encoded by firing rate (i.e. the spike count over a relatively long time window) or by the temporal patterning of spikes [2]–[10]. In temporal coding, information is carried by groups of neurons that fire more or less *synchronously*, as in synfire chains [10], [11]. In rate coding, neuronal firing ideally remains asynchronous across neurons[12-14]. The reliable propagation of synchronous spikes (temporal code) is well understood and relatively easy to implement in computer models [10, 14, 15]. In contrast, the reliable propagation of rate-modulated asynchronous spiking (rate code) is poorly understood and remains difficult to implement in computer models [12]. Indeed, spikes may synchronize as the signal progresses through deeper layers or spike rate may tend toward an attractor state representing quiescence or a fixed rate. In all of the scenarios, rate-based coding is compromised.

Several studies have addressed the conditions required for spike-rate propagation [1], [12]–[16]. Shadlen and Newsome [14] demonstrated the feasibility of rate transmission using leaky integrate and fire (LIF) models receiving balanced excitatory and inhibitory inputs. But Litvak et al [12] showed, using the same parameters quoted by [14], that rate was not reliably transmitted when >2 layers were considered. They concluded that rate transmission in FFNs is highly unlikely. Van Rossum et al [15] showed the feasibility of reliable transmission of instantaneous firing rate (asynchronous spikes) in un-balanced FFNs where the input to each layer is delivered as an injected current, which is not biologically realistic. Kumar et al [13] studied conditions for propagating synchronous spiking and asynchronous firing rate using more complicated and biologically realistic network models. They showed that the coexistence of firing rate and synchrony propagation can be achieved under precise combinations of synaptic strength and connection probability [1]. More recently, Cortes and Vreeswijk [17] showed that pulvinar thalamic nucleus allows for asynchronous spike propagation through the cortex; they supply the input-output firing rate relationship between two cortical areas without manipulating synaptic strengths. It is to be noted that in all previous studies (except Van Rossum et al [15]) the input signal to FFNs is uncorrelated, meaning that the average firing rate is the sole information to be transmitted. More considerations should be given to a correlated (shared) input signal where information of the time-varying firing rate reflected by slow-modulation of asynchronous spikes are to be transmitted. Nevertheless, reliable propagation of the time-varying firing rate under biologically realistic assumptions is rarely considered in the literature.

In this paper, we investigate the *necessary conditions* for reliable propagation of time-varying firing rate –slowly time-varying asynchronous spikes– through FFNs. As reliable transmission of firing rate across several layers of a FFN is achievable if the asynchronous spikes can be maintained almost unchanged in the preceding layers, we explore those conditions in a FFN with only one layer. To this end, we create a FFN composed of excitatory neurons, modeled by leaky integrate and fire (LIF) model, receiving shared input from the previous layer plus background synaptic noise (see **Figure 1**). The novelty of this work lies in (**i**) the development of reduced network model that allows systematic understanding of information propagation in FFNs, and (**ii**) the exploration of necessary conditions for reliable propagation of time-varying firing rates. In the proposed reduced model, synaptic weights are introduced as vectors (rather than as matrices, see **Figure 1**) to encode the common input signal. This enables the use of convex optimization techniques to calculate synaptic weights for maximum information transmission. By incorporating optimum synaptic weights, we show that the network size and the level of background synaptic noise are critical factors for reliable propagation of the time-varying firing rate. The organization of this paper is as follows. In Section II, the reduced network model of a FFN is developed. The constrained and un-constrained optimization methods for calculating synaptic weights of the reduced model are presented in Section III. Necessary conditions for reliable representation of asynchronous spikes are studied in Section IV. And finally, concluding remarks and future directions are provided in Section V.

**Figure. 1:**
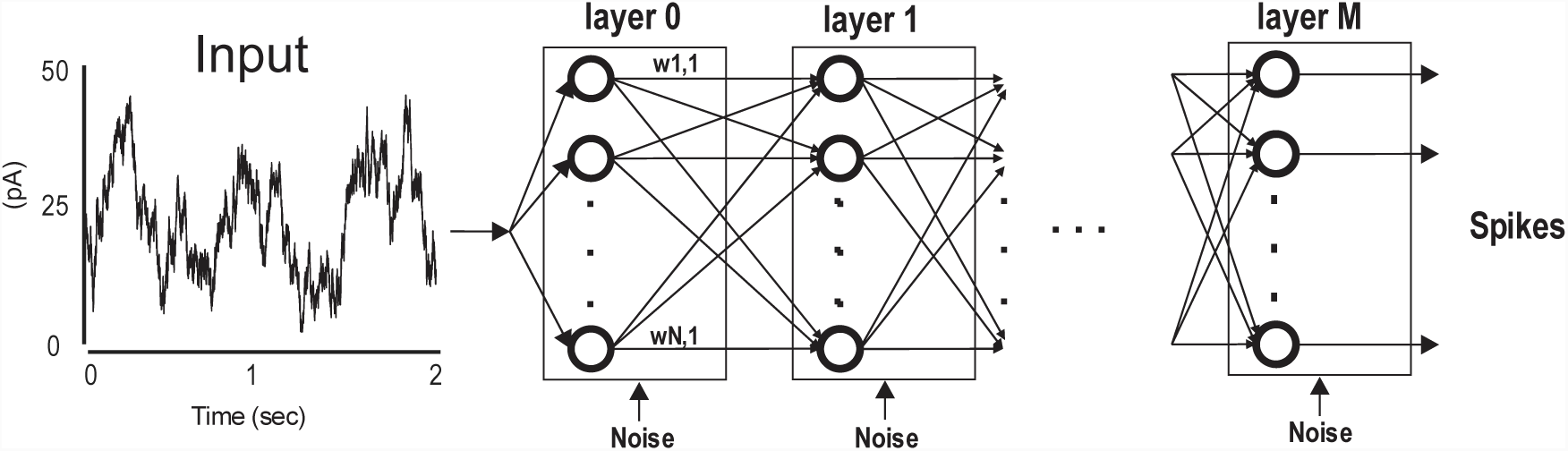
FFN model of information propagation. Schematic representation of information propagation in a feed-forward neural network with all-to-all connectivity comprising N neurons and M layers.

## II. Reduced Network Model of FFN

In this section, the reduced network model of a FFN is derived from its full network model. **Figure. 1** shows a schematic representation of an all-to-all connected multi-layered FFN comprising N neurons and M layers. Each neuron – modeled by LIF model (see Methods) – receives an independent background synaptic noise – modeled by Ornstein-Uhlenbeck (OU) process (see Methods). Neurons in the first layer receive a common slow signal indicating the projection of slowly-modulated asynchronous spikes generated by upstream neurons. Our objective is to find the necessary conditions for reliable information transmission, meaning that the common input can be reconstructed from the spikes of each layer of the FFN. In this paper, we study these conditions for representation of information in the first layer of the FFN.

Although the focus of this study is on the quality of signal processing in the first layer of a FFN where the common input signal is generated by an OU process (time constant = 50 msec, see Methods), the problem of reduced network model is stated for more general scenarios. In fact, we consider that the common input of layer 1 is composed by spikes in layer 0 (see **Figure. 1**), and this problem statement can be generalized to any consecutive layers in a multi-layered FFN.

For each neuron in the first layer of the FFN, the post-synaptic potential (input, *I*) is produced by passing (convolving) the spikes of the preceding layer (layer 0), in all-to-all connection case, through an identical synaptic waveform. Then, the spikes of each neuron is generated by feeding the weighted sum of those post-synaptic potentials into the LIF model (note that each neuron receives an independent background noise as well). The spike train of neuron *i* in the first layer,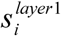, can be written as follows.

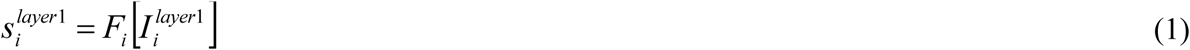

where *I*_*i*_ and *F*_*i*_ indicate the (synaptic) input and the model neuron (LIF, see Methods) of neuron *i*. As neurons, in this paper, are homogeneous, *F*_*i*_ *= F* for all *i* = 1, …, N. And, 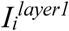 can be expressed by:

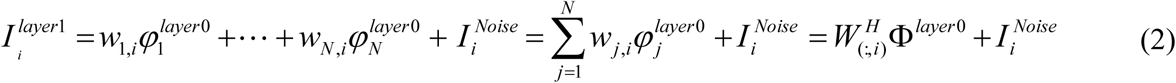

where 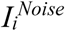 stands for background synaptic noise received by neuron *i*, and modeled by Gaussian noise with zero mean and standard deviation *σ,* i.e., the level of background synaptic noise.

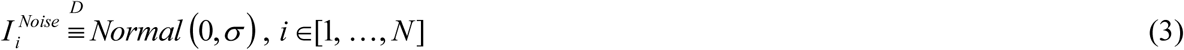

In (2), *W* represents the synaptic weight matrix.

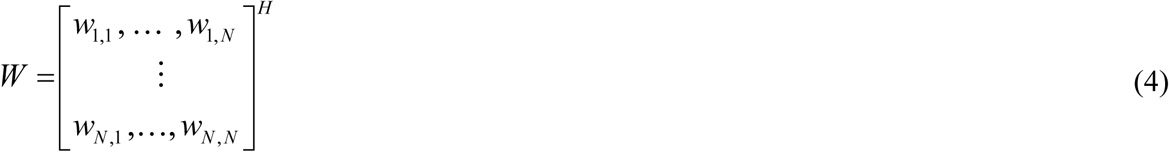

And, Φ^*layer*0^ (non-weighted post-synaptic potentials) indicates a vector comprising spikes (layer 0) convolved with the synaptic waveform, and it can be expressed as follow.

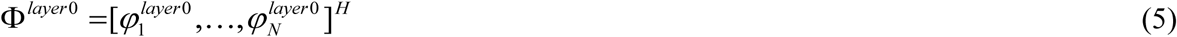

where,

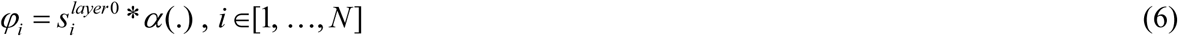

*α* (.) is an identical synaptic waveform modeled by a double exponential function of *τ*_*rise*_ = 0.5 msec & *τ*_*fall*_ = 3 msec, and ‘*’ stands for the convolution function.

Here, we define the objective function underlying maximum information representation in the firs layer of a FFN, that is equivalent to minimization of the l2 norm between the instantaneous firing rate in the first layer and that in the initial layer (layer 0 in **Figure. 1**).

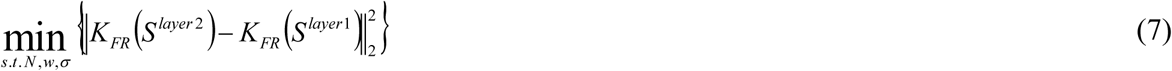

where, K_FR_(S) = S * *Kn*, represents the instantaneous firing rate, and *Kn* is a Gaussian kernel with standard deviation of 50 ms. 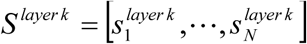 shows the vector of all neuron’s spikes, and k indicates the layer index. Since the exact spike timing cannot be reproduced in even purely information propagation, we consider the instantaneous firing rate with a moderate time window of 50 ms – equivalent to time constant of the input – to compare the spikes in the initial and first layers.

### Network Model Reduction

Figure. 2 shows a reduced network model which is equivalent to the full model shown in Figure. 1 under some biologically-reasonable assumptions, namely, homogeneity of neurons and synapses, as well as the convergence of upstream neurons that produce a common input to the neurons in FFN.

**Figure. 2:**
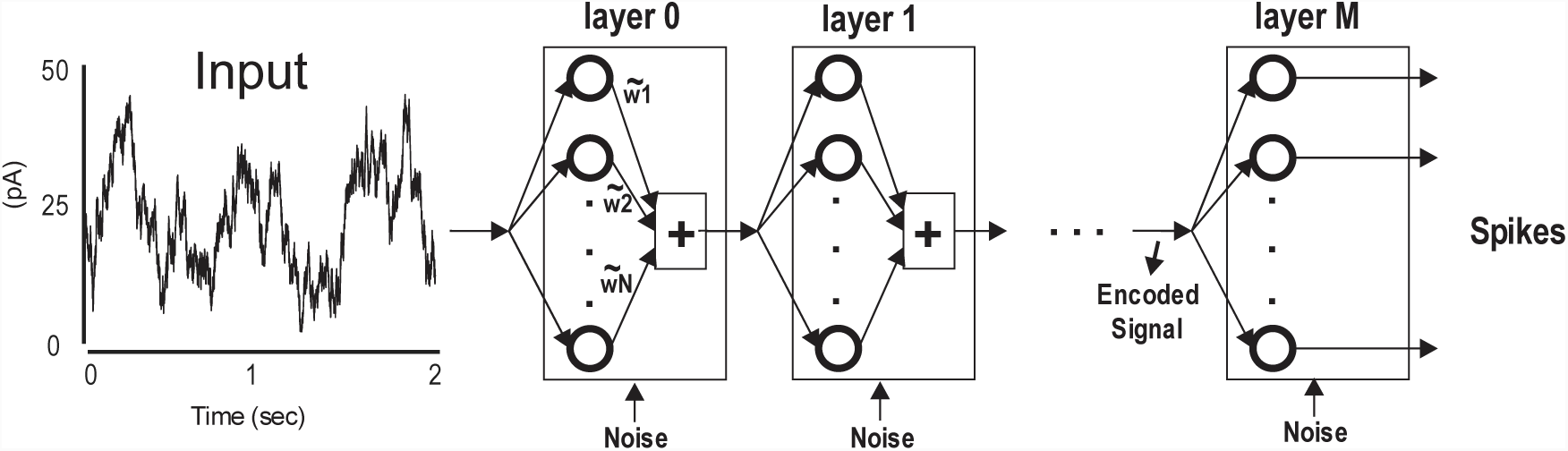
Schematic representation of an abstract model of propagation equivalent to Figure 1.

Given abovementioned assumptions, one can express the common input to the neurons of an arbitrary layer as:

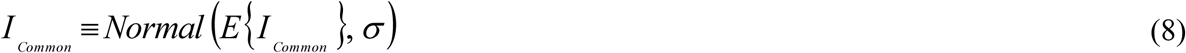

where E{.} is the expectation function, and *σ* is the level of background synaptic noise. One can see that the common input has a Gaussian distribution whose variability is produced by the synaptic noise that each neuron (homogeneous) receives, that is *I*_*Common*_= *E*{*I*_*Common*_}+ *Normal* (0,*σ)* (see (2)). Considering the homogeneity of each neuron, the mean of the common input is equal to that of each individual neuron, that is

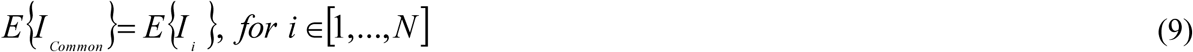

From (2), we have,

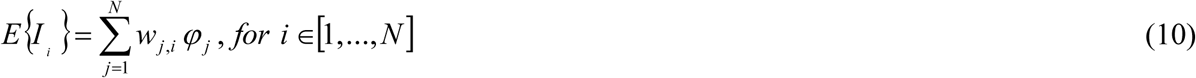

In fact, (10) implies that the mean of the input signal to all neurons in a specific layer is identical. Thus, we can write,

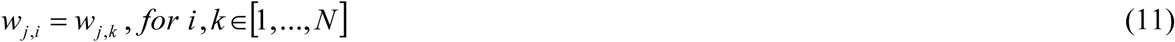

Thus, (11) offers a network model reduction, as shown in **Figure. 2**, with respect to abovementioned assumptions. Given (9), (10) and (11), one can replace the synaptic matrix W with the synaptic vector 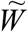, where,

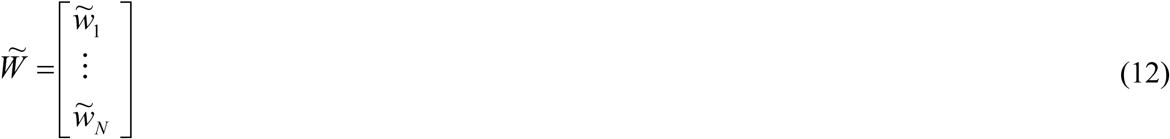

the mean of the input to a neuron, i.e., identical for all neurons in a specific layer, can be written as:

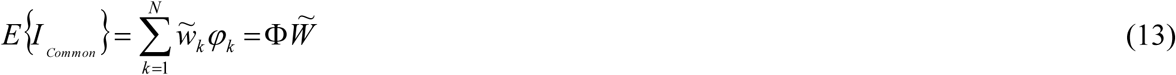

Therefore, unlike *W* that reflects the synaptic weights, 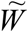 expresses the contribution of neurons in constructing (decoding) the input signal, i.e., each neuron has a certain contributing weight which is identically distributed to all following neurons.

## III. A Tractable Optimization Framework to Calculate Synaptic Weights

The reduced network model implies that the mean of common input to each layer of a FFN can be described by a matrix of post-synaptic potentials, Φ, in the preceding layer multiplied by the vector of synaptic weights (see (13)). Thus, the generated spikes by a neuron in the first layer, as expressed in (1), can be rewritten as follows.

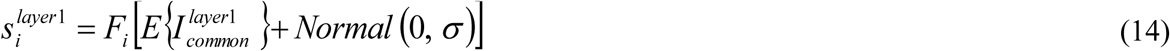

And equivalently, the spikes matrix of all neuron in the first layer will be expressed by:

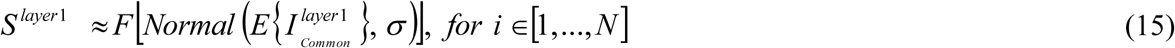

In fact, the objective function (7) is equal to:

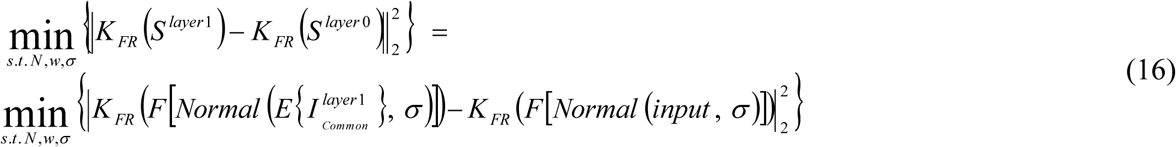

Here we use a rational to simplify (16), and solve it in an indirect but innovative way. Minimization of (16) implies that the reconstructed input by spikes of neurons in the first layer of the FFN should be equal (ideal case) to the original input (in layer 0). This interpretation is equivalent to maximization of information representation from decoding perspective [2]. Using this interpretation, we write a new objective function as below.

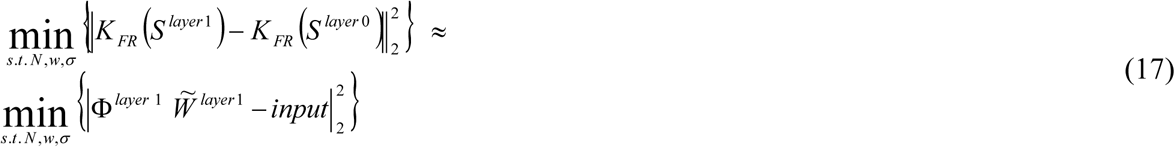

As we aim to calculate the synaptic weights through a tractable convex optimization problem, we write (17), given *N* and *σ*, as follow.

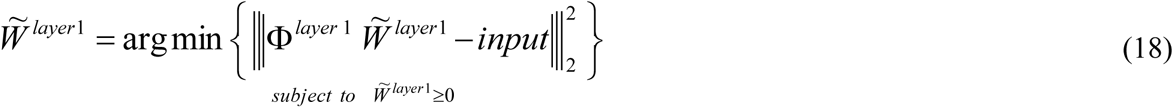

We have derived constrained and unconstrained optimization solutions for (18). We use constrained optimization technique, “*lsqnonneg*” in MATLAB. As well, we use unconstrained optimization (Least-square method) where the negative weights (after calculation) are set to zero. In this case, we have:

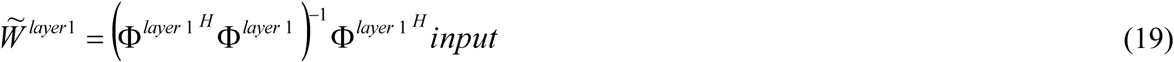

where *H* indicates a matrix Hessian.

## IV. Necessary Conditions for Reliable Representation of Asynchronous Spikes

We study, in this section, the effect of synaptic weights 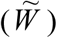, network size (*N*), and level of background synaptic noise (*σ*) on the reliability of information (asynchronous spikes) representation in the first layer of a FFN. The reduced network model and corresponding optimization solutions are used in the following numerical simulations.

**Figure. 3** shows the main steps underlying the numerical simulations for a FFN comprising 200 neurons. Each neuron receives an independent background synaptic noise with the standard deviation of 15 pA. The common input in layer 0 is produced by OU process with an autocorrelation time of 50 msec. Figure. 3(A) shows the input (inset: distribution of amplitudes) and generated spikes (raster plot) of neurons in the first layer of the FFN (inset: firing rate of all individual neurons). The reconstructed inputs using constrained and unconstrained optimization techniques as well as the distribution of synaptic weights are shown in Figure. 3 (B) and (C), respectively. In this example, the synaptic weights (and accordingly the reconstructed inputs) are almost the same using both unconstrained and constrained techniques. Note that the dimension of synaptic weights is expressed by pA/mV based on (19) and the excitatory synapses (excitatory reversal potential = 0 mV).

**Figure. 3:**
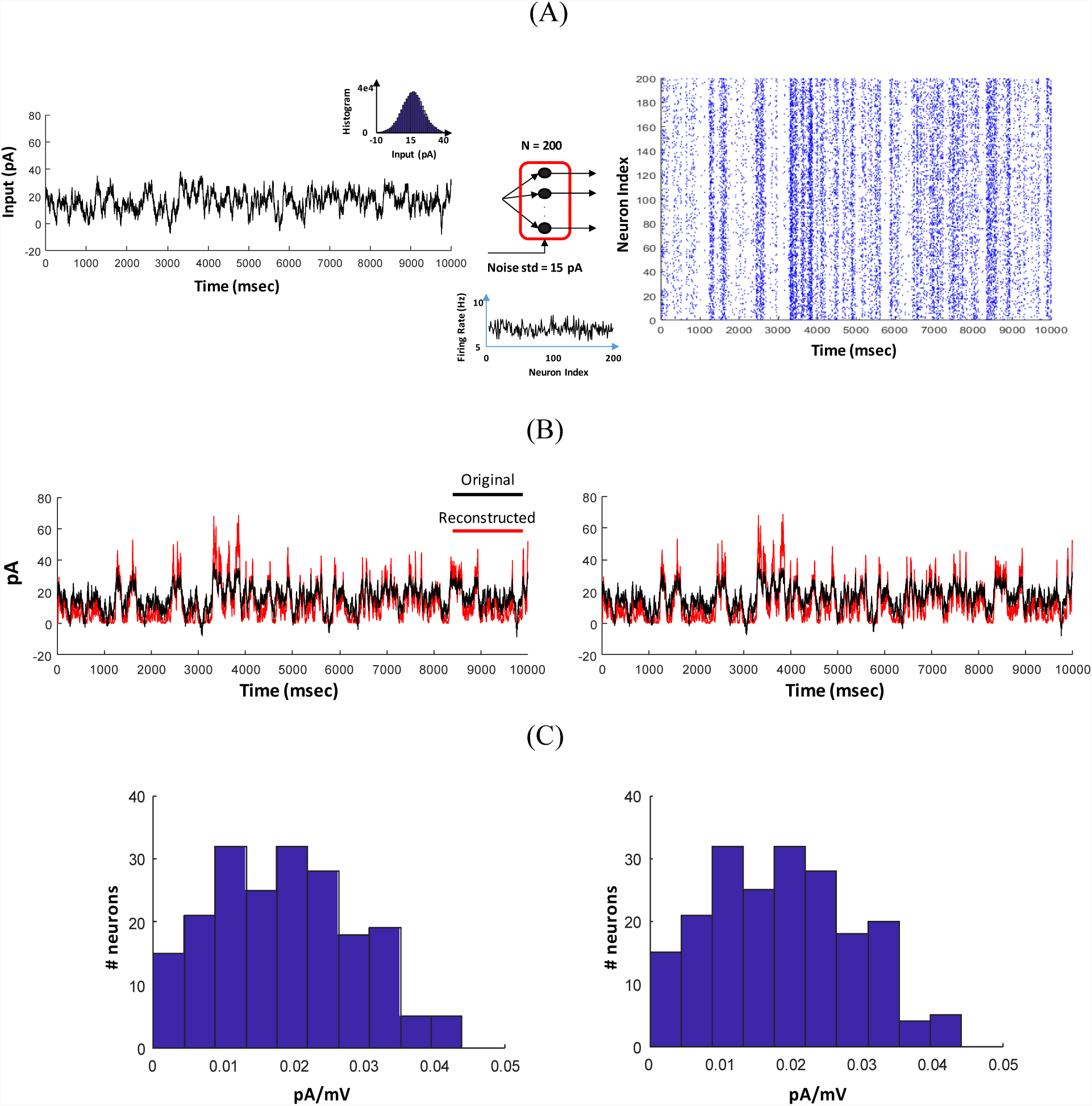
Reconstruction of a common slow input signal from the generated spikes of the first layer of a FFN using constrained and unconstrained optimization techniques. (A) common slow signal (left) and generated spikes (N = 200 & std of noise = 15 pA). Insets of (A): statistical characteristics of input (OU process of time constant 50 msec) and spikes (average firing rate = 7 Hz). (B) Original vs. reconstructed input signals using constrained (left) and unconstrained (right) optimization techniques. (C) The histogram of connectivity weights (corresponding to the abstract model of propagation) are shown at the bottom of each reconstructed signal in (B).

### Network Size has a key role to preserve information of the preceding layer

Using the same procedures as explained in **Figure. 3**, we reconstruct the input signal of a FFN with different network sizes. **Figure. 4** indicates that the reconstructed input for N = 500 is better representing the original input than that for N = 50. The level of background synaptic noise is 15 pA for both scenarios. The distribution of synaptic weights is also different for different network sizes. For N = 500, almost 150 neurons have very weak contributions in signal reconstruction (transmission), implying that an optimum network size (no redundant weights) might exist.

**Figure. 4:**
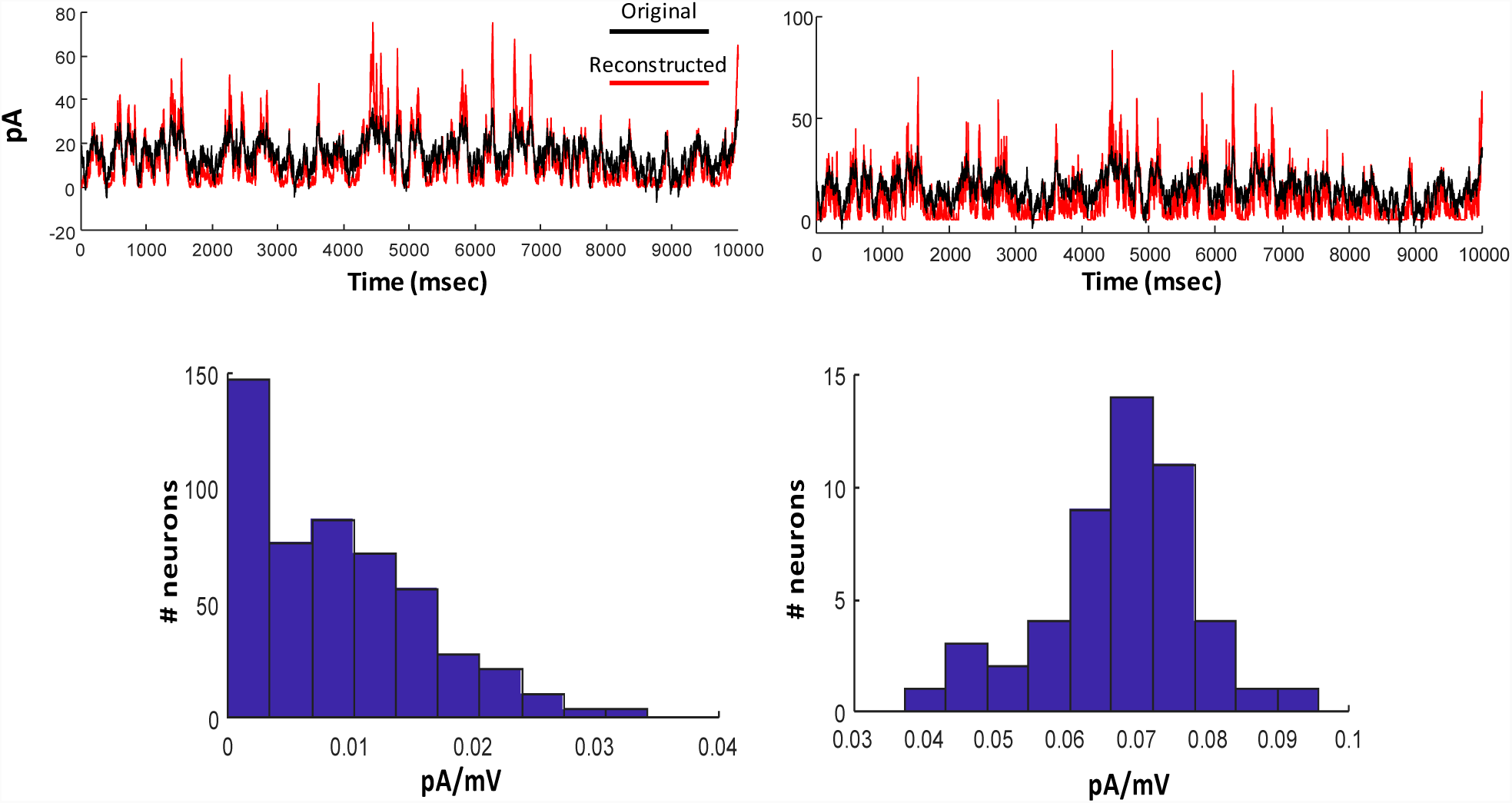
Original vs. reconstructed signals using unconstrained optimization of a FFN with N = 500 (left) and 50 (right) for noise std = 15 pA. The histogram of connectivity weights (corresponding to the abstract model of propagation) are shown at the bottom of each reconstructed signal.

To explore whether an optimum network size exists, the mean squared error between the original and reconstructed inputs are calculated for different network sizes given a consistent synaptic noise (*σ* = 15 pA). **Figure. 5** shows this error for different network sizes. As expected, the error has a minimum for N = 200, i.e., information (asynchronous spikes) transmission, given a fixed level of synaptic noise, is maximized for a certain network size.

**Figure. 5:**
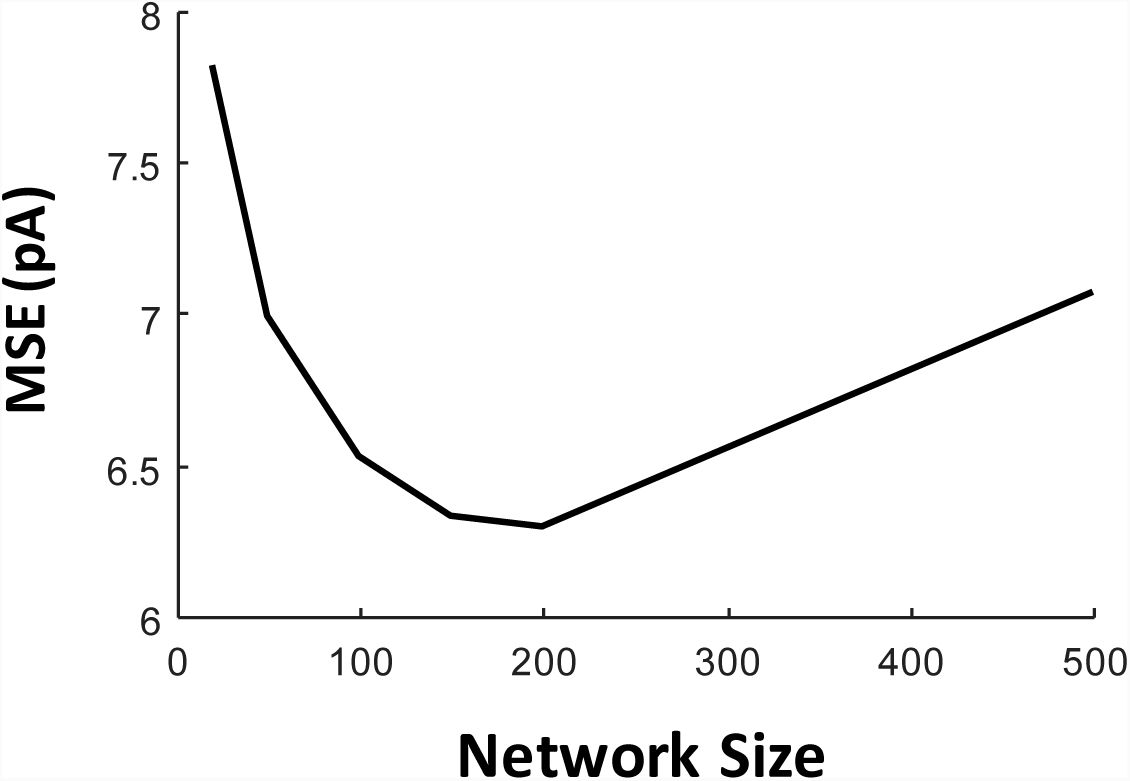
Quality of reconstructed signal depends on network size. Error plot of information representation for different network sizes for noise std = 15 pA.

### Quality of reconstructed signal depends on the level of background synaptic noise

In order to study whether the quality of reconstructed signals in the first layer of a FFN depends on the level of background synaptic noise, we calculate the error shown in **Figure. 5** for a higher level of noise, *σ* = 25 pA. As can be seen in **Figure. 6**, the error decreases when network size increases. In fact, **Figures. 5-6** demonstrate that the reliability of information transmission depends on both network size and the level of background synaptic noise. It is to be noted that the noise std of 25 pA is equivalent to the level of background synaptic noise observed in-vivo [18]. Other studies based on reverse correlation analysis [19] showed that this level of noise maximizes coding fraction of asynchronous spikes in reconstruction of slow signals.

**Figure. 6:**
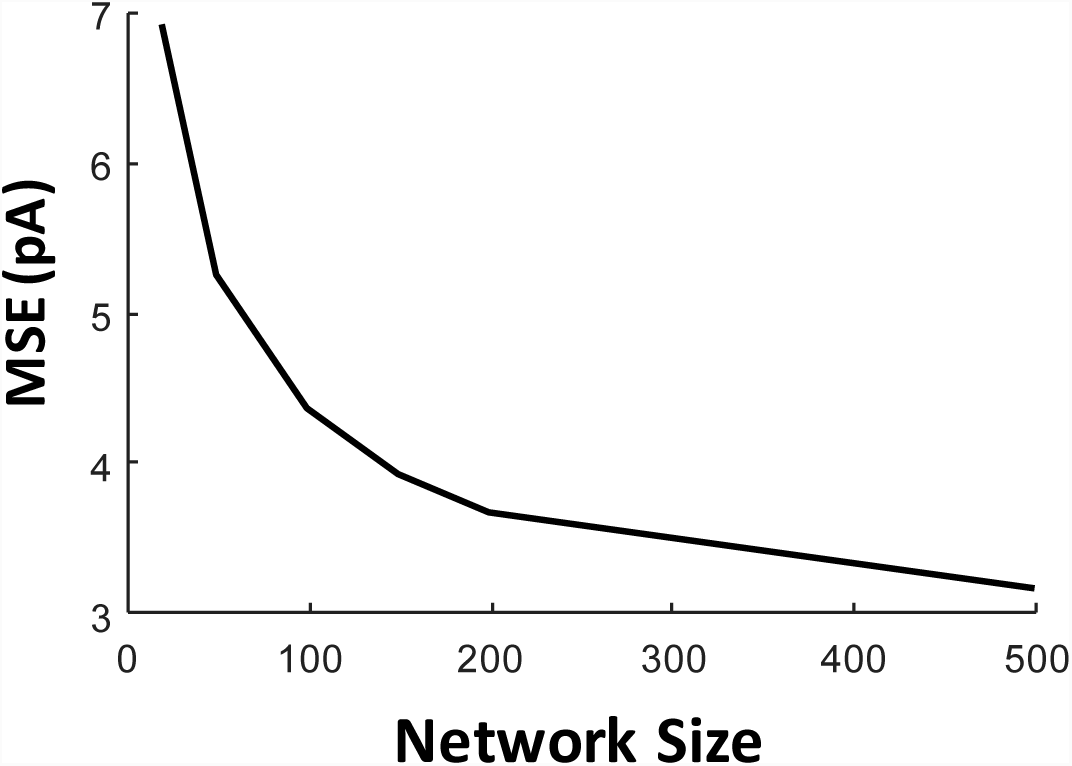
Quality of reconstructed signal depends on the level of background noise. Error plot of information representation for different network sizes, and a fixed noise level (std = 25 pA).

To better visualize the quality of information representation for different levels of background synaptic noise, the input signal is reconstructed by FFNs with fixed network sizes (N=200) and different levels of noise, *σ* = 15 pA and *σ* = 25 pA. **Figure. 7** shows that the reconstructed input by a FFN with *σ* = 25 pA is better – compared to *σ* = 15 pA – describing the original input. Moreover, unlike the FFN with *σ* = 15 pA, the distribution of synaptic weights of the FFN with *σ* = 25 pA does not comprise any weak synapses (redundant neurons).

**Figure. 7:**
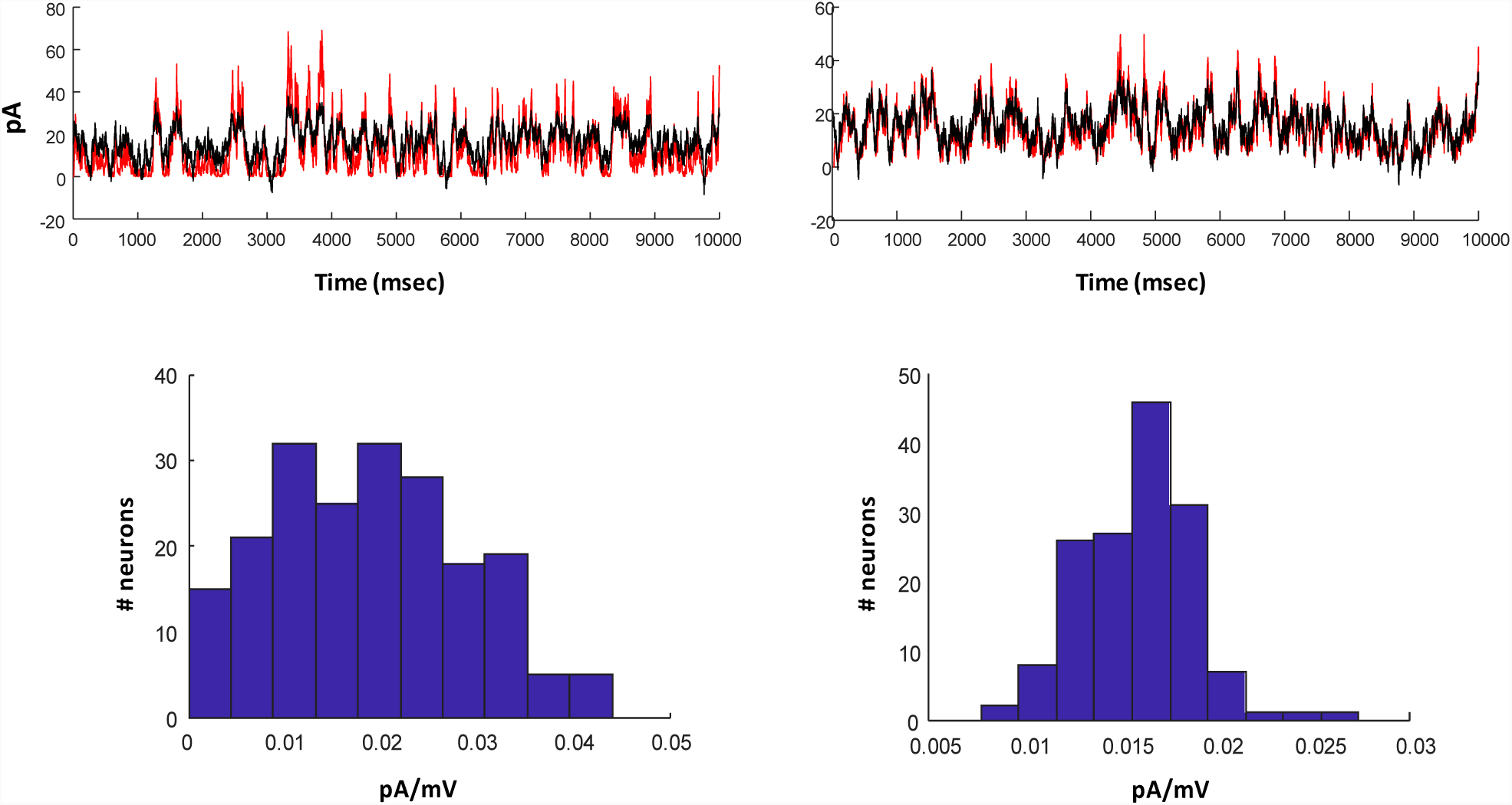
An example of reconstructed signal for N = 200 and different level of background synaptic noise, noise std = 15 pA (left) and noise std = 25 pA (right). The histogram of connectivity weights (corresponding to the abstract model of propagation) are shown at the bottom of each reconstructed signal.

Although the error decreases for larger network sizes in **Figure. 6**, it should be regulated by biologically realistic levels of synaptic weights. In fact, the amplitude of the synaptic waveform indicates how much membrane potential of a neuron (in the next layer) depolarizes (for excitatory neurons) given pre-synaptic spikes. The values of estimated synaptic weights using our optimization framework should be rescaled accordingly, and biologically unrealistic synaptic weights should be excluded.

## V. Conclusion

Necessary conditions for reliable information propagation in a multi-layered FFN were investigated in this paper. Previous studied have addressed those conditions for transmission of *synchronous spikes* as well as the *mean of firing rate* in multi-layered FFNs. However, these conditions barely remain valid for transmission of *asynchronous spikes* – slowly time-varying firing rate. In this paper, we investigated the necessary conditions for reliable transmission of asynchronous spikes in a FFN. Specifically, we explored those conditions within a single layered FFN where the quality of represented information can be better controlled with respect to certain parameters (conditions). One can agree that the instantaneous firing rate of spikes in two consecutive layers of a FFN – with all-to-all connectivity comprising homogeneous neurons and synapses – should be almost similar if their underlying common input are the same. This was our rationale to derive an objective function that is a L2-norm error between the original input and that reconstructed by spikes of the first layer of the FFN. After establishing that a FFN with all-to-all connectivity can be represented by an abstract network model (i.e., vector representation of synaptic weights), we used optimized synaptic weights, which minimizes the objective function (equivalent to maximizing coding fraction [2]), to study the optimal conditions for reliable representation of asynchronous spikes. We found that not only the values of synaptic weights but also two other factors, namely (i) the network size and (ii) the level of background synaptic noise are critical to enable reliable information representation.

Validation of these conditions in reliable transmission of asynchronous spikes in multi-layered FFN creates our future lines of research. It is to be noted that other biological factors like heterogeneous synapse and dendritic integration can significantly change the quality of signal representation (coding fraction). we will study the effect of these factors in our future studies.

## Acknowledgement

We thank Dr. Steve Prescott for his insight and comments on the original version of this study.

## Methods

### LIF model

Neurons were modeled as follows.

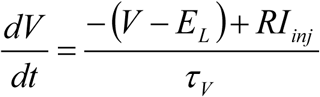

where *E*_L_=−70 mV, *R* = 1 MΩ, and *τ*_*V*_ = 10 msec. *I*_*inj*_ indicates the injected current (slow signal in this paper). Spike occurs when V≥Vth, where *V*_*th*_ = −40 mV and the reset voltage is −90 mV.

### Slow Signal and background synaptic noise

The slow signal and background synaptic noise are generated using an Ornestein-Uhlenbeck (OU) process of autocorrelation time of 50 msec and 5 msec, respectively. The mean and variance of the slow signal (in layer 0) is 16 pA and 6 pA, respectively.

## References

[1] A. Kumar, S. Rotter, and A. Aertsen, “Spiking activity propagation in neuronal networks: reconciling different perspectives on neural coding,” Nat. Rev. Neurosci., vol. 11, no. 9, pp. 615–627, Sep. 2010.

[2] B. & Noble, “Spikes: Exploring the Neural Code / Edition 1,” Barnes & Noble. [Online]. Available: https://www.barnesandnoble.com/w/spikes-fred-rieke/1117355212. [Accessed: 01-Feb-2019].

[3] S. Panzeri, R. S. Petersen, S. R. Schultz, M. Lebedev, and M. E. Diamond, “The Role of Spike Timing in the Coding of Stimulus Location in Rat Somatosensory Cortex,” Neuron, vol. 29, no. 3, pp. 769–777, Mar. 2001.

[4] M. A. Montemurro et al., “Role of Precise Spike Timing in Coding of Dynamic Vibrissa Stimuli in Somatosensory Thalamus,” J. Neurophysiol., vol. 98, no. 4, pp. 1871–1882, Oct. 2007.

[5] Y. Zuo, H. Safaai, G. Notaro, A. Mazzoni, S. Panzeri, and M. E. Diamond, “Complementary Contributions of Spike Timing and Spike Rate to Perceptual Decisions in Rat S1 and S2 Cortex,” Curr. Biol., vol. 25, no. 3, pp. 357–363, Feb. 2015.

[6] M. London, A. Roth, L. Beeren, M. Häusser, and P. E. Latham, “Sensitivity to perturbations *in vivo* implies high noise and suggests rate coding in cortex,” Nature, vol. 466, no. 7302, pp. 123–127, Jul. 2010.

[7] C. A. Runyan, E. Piasini, S. Panzeri, and C. D. Harvey, “Distinct timescales of population coding across cortex,” Nature, vol. 548, no. 7665, pp. 92–96, Aug. 2017.

[8] S. Panzeri, C. D. Harvey, E. Piasini, P. E. Latham, and T. Fellin, “Cracking the Neural Code for Sensory Perception by Combining Statistics, Intervention, and Behavior,” Neuron, vol. 93, no. 3, pp. 491–507, Feb. 2017.

[9] J. Kremkow, A. Aertsen, and A. Kumar, “Gating of Signal Propagation in Spiking Neural Networks by Balanced and Correlated Excitation and Inhibition,” J. Neurosci., vol. 30, no. 47, pp. 15760–15768, Nov. 2010.

[10] M. Abeles, Y. Prut, H. Bergman, and E. Vaadia, “Synchronization in neuronal transmission and its importance for information processing,” in Progress in Brain Research, vol. 102, J. Van Pelt, M. A. Corner, H. B. M. Uylings, and F. H. Lopes Da Silva, Eds. Elsevier, 1994, pp. 395–404.

[11] M. Diesmann, M.-O. Gewaltig, and A. Aertsen, “Stable propagation of synchronous spiking in cortical neural networks,” Nature, vol. 402, no. 6761, pp. 529–533, Dec. 1999.

[12] V. Litvak, H. Sompolinsky, I. Segev, and M. Abeles, “On the Transmission of Rate Code in Long Feedforward Networks with Excitatory–Inhibitory Balance,” J. Neurosci., vol. 23, no. 7, pp. 3006–3015, Apr. 2003.

[13] A. Kumar, S. Rotter, and A. Aertsen, “Conditions for Propagating Synchronous Spiking and Asynchronous Firing Rates in a Cortical Network Model,” J. Neurosci., vol. 28, no. 20, pp. 5268–5280, May 2008.

[14] M. N. Shadlen and W. T. Newsome, “Noise, neural codes and cortical organization,” Curr. Opin. Neurobiol., vol. 4, no. 4, pp. 569–579, Aug. 1994.

[15] M. C. W. van Rossum, G. G. Turrigiano, and S. B. Nelson, “Fast Propagation of Firing Rates through Layered Networks of Noisy Neurons,” J. Neurosci., vol. 22, no. 5, pp. 1956–1966, Mar. 2002.

[16] S. Wang, W. Wang, and F. Liu, “Propagation of Firing Rate in a Feed-Forward Neuronal Network,” Phys. Rev. Lett., vol. 96, no. 1, p. 018103, Jan. 2006.

[17] N. Cortes and C. van Vreeswijk, “Pulvinar thalamic nucleus allows for asynchronous spike propagation through the cortex,” Front. Comput. Neurosci., vol. 9, 2015.

[18] A. Destexhe, M. Rudolph, J.-M. Fellous, and T. J. Sejnowski, “Fluctuating synaptic conductances recreate in vivo-like activity in neocortical neurons,” Neuroscience, vol. 107, no. 1, pp. 13–24, Nov. 2001.

[19] M. Lankarany and S. A. Prescott, “Multiplexed coding through synchronous and asynchronous spiking,” BMC Neurosci., vol. 16, no. Suppl 1, p. P198, Dec. 2015.

